# Diet composition of introduced Barn Owls (*Tyto alba javanica*) in urban area in comparison with agriculture settings

**DOI:** 10.1101/574277

**Authors:** Safwan Saufi, Shakinah Ravindran, Noor Hisham Hamid, Cik Mohd Rizuan Zainal Abidin, Hamdan Ahmad, Abu Hassan Ahmad, Hasber Salim

**Author notes:** Corresponding author (HS). These authors contributed equally to the work (Safwan Saufi, Shakinah Ravindran, Hasber Salim). These authors also contributed equally to the work (Noor Hisham Hamid, Cik Mohd Rizuan Zainal Abidin, Hamdan Ahmad, Abu Hassan Ahmad).

## Abstract

This study investigated the diet of introduced barn owls (*Tyto alba javanica*, Gmelin) in the urban area of the Main Campus of Universiti Sains Malaysia, Penang, Malaysia, based on collected regurgitated pellets. We also compared the diet of introduced urban barn owls with the diet of barn owls from two agricultural areas, i.e. oil palm plantations and rice fields. Pellet analysis of barn owls introduced in the urban area showed that commensal Norway rats, *Rattus norvegicus*, made up the highest proportion of the diet (65.37% prey biomass) while common shrews, *Suncus murinus* were the second highest consumed prey (30.12% prey biomass). Common plantain squirrel, *Callosciurus notatus*, made up 4.45% of the diet while insects were taken in a relatively small amount (0.046% prey biomass). Introduced barn owls showed a preference for medium-sized prey, i.e. 40 to 120g (52.96% biomass and 38.71% total). In agricultural areas, *Rattus argentiventer* predominated the diet of barn owls (98.24% prey biomass) in rice fields while Malayan wood rats, *Rattus tiomanicus*, were the most consumed prey in oil palm plantations (99.5% prey biomass). Food niche breadth value was highest for barn owls introduced in an urban area with a value of 2.90, and 1.06 in rice fields and 1.22 in oil palm plantations. Our analysis reiterates the prey preference of barn owls in various landscapes for small mammals. Our results also indicate the suitability of utilizing barn owls as a biological control not only in agricultural areas, but also as a biological control agent for commensal rodent pests in urban areas.

## Introduction

The barn owl, *Tyto alba* (Tytonidae), is a common species of owls which occurs on almost all continents and in most open lands and farmlands (Bunn et al., 1982; Taylor, 1994). Like many other cosmopolitan nocturnal raptors, barn owls display an astonishing breadth of habitat association and have been able to adapt and persist in areas that are becoming urbanized (Hindmarch et al., 2017). The diet of barn owls has been well studied throughout its range because of the ease of identifying prey remnants recovered inside regurgitated pellets. Owls swallow their prey whole and expel pellets, which are composed of undigested remains such as bones, compacted in hair and feathers (Taylor, 1994). Analysis of barn owl pellets have provided information on the diet composition of owls and dynamics of prey species communities within the owl foraging areas (Alivizatos & Goutner, 1999; Kitowski, 2013).

The diet of barn owls in agricultural areas has been extensively studied throughout its range (Jaksic et al., 1982; Marti, 2010; Paspali et al., 2013). In most part of their foraging range, barn owls feed primarily on small mammals, i.e. rats, mice, voles and shrews, with birds, insects, amphibians, reptiles and invertebrates taken in relatively smaller amounts (Bunn et al., 1982). In Peninsular Malaysia, several studies on the food selection of barn owl in major agricultural crop areas report rats as the major prey. Diet analysis of the owl’s regurgitated pellets show that rats comprise more than 98% of the prey in oil palm plantations (Lenton, 1984) and 94.7% in rice fields (Hafidzi et al., 1999).

Its renowned role as an efficient small mammal predator has led to barn owls being introduced in various landscapes. Barn owls have been introduced in islands (Au & Swedberg, 1966; Emmerson & Ascani, 1985), agricultural areas (Hafidzi & Naim, 2003b; Rizuan et al., 2017) and semi-urban areas (Meyer, 2008) for the purpose of controlling pest rodent populations. Barn owls are also translocated as part of reintroduction programs for declining local barn owl populations (Meek et al., 2003). In this study, Southeast Asian barn owls, *Tyto alba javanica*, were translocated from their native agricultural habitats and introduced to the urban-garden area of the Main Campus of Universiti Sains Malaysia to serve as a biological control agent against the rat pest population. Here, we report the analysis of the diet composition of introduced barn owls in an urban area, and compared the diet of introduced urban barn owls to the diet of barn owls in oil palm plantations and rice fields in Peninsular Malaysia.

## Materials and methods

This study was carried out in strict accordance with the recommendations in the Animal Research and Service Centre, Universiti Sains Malaysia (USM). The study protocol was approved by the Animal Ethics Committee USM (Approval number: USM / Animal Ethics Approval / 2015 / (96) / (629)). Permit for the study was approved by the Department of Wildlife and National Parks, Peninsular Malaysia (Permit number: JPHL&TN(IP): 60-4/1/13 Jilid 20 (28)).

### Study area and introduced barn owls

In total, 24 barn owls were released intermittently from April 2016 to August 2018 in the urban-garden area of the Main Campus of Universiti Sains Malaysia (USM), Penang, Malaysia (5.3579° N, 100.2943° E). Prior to the release of barn owls, 14 artificial nest boxes were set up around on the campus area. Providing nest boxes is a common practice to attract barn owls and increase nesting performance and hence sustain barn owl population. Two types of artificial nest boxes, i.e. wooden and fibreglass, were installed early in January 2016 and scattered around the campus at open areas of vegetation (Figure 1).

**Figure 1:**
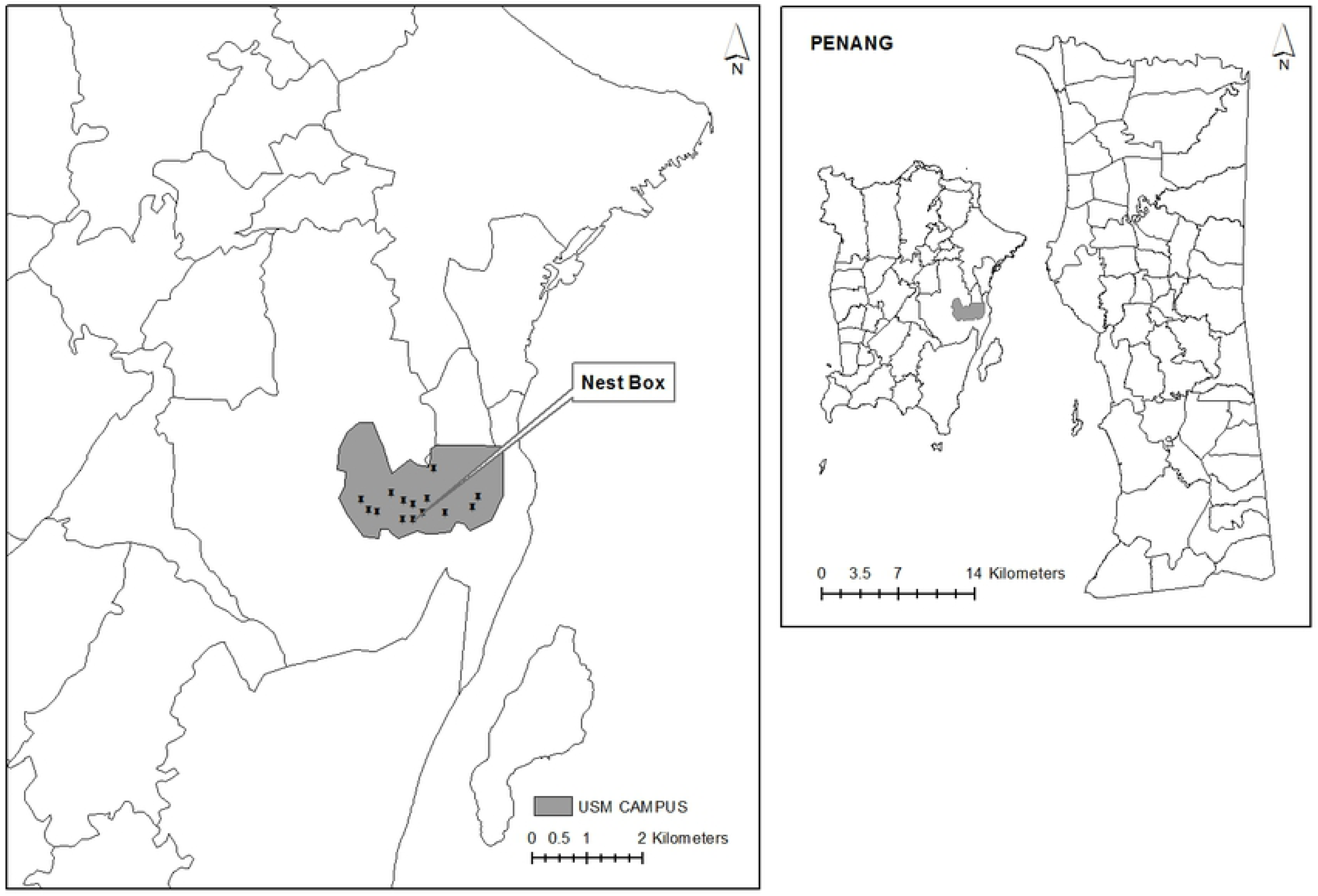
Study site of introduced barn owl and location of nest boxes.

The translocated barn owls were harvested from three different locations in Peninsular Malaysia; oil palm plantations at the Tun Razak Agricultural Research Centre, Bandar Jengka Pahang (3.777967° N, 102.517238° E), rice fields of Bumbung Lima, Kepala Batas, Pulau Pinang (5.51707° N, 100.4265° E) and rice fields in the Kerian District, Parit Buntar, Perak (5.0081° N, 100.5394° E). The owls were temporarily held in the USM Aviary (5.35791944° N, 100.29416667° E) for about one month before release to allow the birds to acclimatize to their new urban surroundings.

All introduced barn owls were banded with customized metal leg bands prior to release. Transmitters were fitted to the owls using backpack style (Saufi et al., 2018). The transmitter and harness weighed approximately 9 g, i.e. less than 2 % of total body mass of the barn owls (range between 430 g to 580 g) to avoid affecting bird behaviour and movement (Gaunt et al., 1997). VHF-radio telemetry (TRX-48S, Wildlife Materials Inc.) was used to observe the post-release movement of released barn owls. Each owl was followed for at least 10 cumulative days immediately following its release, starting from dusk (2000 hours) to dawn (0630 hours). Radio-tracking was initially done from vehicles and when a signal was detected, tracking was done on foot till the strongest signal could be detected. The last detected location of an owl during a tracking session is crucial as it determines the owl roosting site of the day, from which there is a high probability of finding a regurgitated pellet.

Regurgitated pellets of introduced barn owls were collected from August 2018 to December 2018 at various locations scattered around the campus. Several structures were identified within the campus that were used regularly by barn owls as perching and roosting sites and pellets were collected on the ground below these sites.

### Diet of barn owls in agricultural areas

Barn owls in rice fields were sampled in rice fields of Bagan Serai, Perak and Kepala Batas, Penang. Surveys for barn owl nest boxes and roosting sites were conducted from August 2017 to July 2018. Barn owls in oil palm plantations were sampled from the plantations in Tun Razak Agricultural Research Centre, Bandar Jengka, Pahang. Survey for pellet samples were conducted July 2017 to August 2018. Pellets samples from both agricultural settings were collected in and around nest boxes and identified perching sites of barn owls.

### Pellet analysis

Pellets were soaked in water individually and processed carefully by taking them apart (Terry, 2004). Bone remnants from pellets were preserved in alcohol prior to identification. For rodent identification, skull and lower jaw of the prey were used for identification down to species level following the identification key of Harrison (1962). A scientific calliper (Mitutoyo U.S.A) was used to measure the size of bones to determine the size of the prey (0.01 mm accuracy). If a skull and lower jaw were not present, measurement of the femur and humerus bone was done to distinguish between juveniles and adults. Insects found in pellets were identified using Borror and White (1970) while other vertebrate prey were determined up to family level using identification keys by Beisaw (2013).

The biomass of prey items recovered from pellets were estimated using a standard log-log regression of right mandible length as a function of body weight (Morris, 1979; Hamilton, 1980; Marti, 2009). The food niche breadth (FNB) of barn owls in all the areas was calculated to determine the dietary diversity of barn owls in each habitat. Food niche breadth (FNB) (Levins, 1968) was calculated as follows:

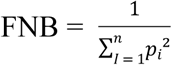

Where *p* is the proportion to prey category *i* in the barn owl diet. Higher values on this index represent a higher diversity of the diet.

## Results

### Diet of barn owls

A total of 252 pellets were collected and 10 groups of animal taxa were identified from prey remnants from all three different study habitats (Table 1). Small mammals from the family Muridae were the staple prey in all three different habitats, though the main prey species differed in each habitat.

**Table 1:** Diet composition of introduced urban barn owls and in agricultural areas.

| Prey species | Urban (%) |  | Rice field (%) |  | Oil palm Plantation (%) |  |
| --- | --- | --- | --- | --- | --- | --- |
|  | Biomass | Individuals | Biomass | Individuals | Biomass | Individuals |
| *R. norvegicus* | 65.37 | 45.05 | 0 | 0 | 0 | 0 |
| *R. argentiventer* | 0 | 0 | 98.24 | 96.77 | 0 | 0 |
| *R. tiomanicus* | 0 | 0 | 0 | 0 | 94.35 | 90.16 |
| House shrew | 30.12 | 35.16 | 1.28 | 2.15 | 0 | 0 |
| Bird | 0 | 0 | 0 | 0 | 0.76 | 1.63 |
| Reptiles | 0 | 0 | 0 | 0 | 0.66 | 0.81 |
| Amphibian | 0 | 0 | 0.46 | 1.07 | 0.47 | 1.63 |
| Grasshopper | 0.11 | 5.49 | 0 | 0 | 0.02 | 3.27 |
| Termites | 0.002 | 12.10 | 0 | 0 | 0 | 0 |
| Plantain squirrel | 4.45 | 2.20 | 0 | 0 | 3.71 | 2.45 |
| FNB | 2.90 |  | 1.06 |  | 1.22 |  |

\*FNB = food niche breadth

A total of 62 individual pellets were collected from barn owls introduced in urban areas and 95.49% of prey biomass of the diet were composed of commensal rodent pests. The Norway rat, *Rattus norvegicus* was the most preyed on; making up 45.05% of total pellet contents and 65.37% of prey biomass. House shrews, *Suncus murinus* were the second highest consumed prey item of barn owls in the urban area (35.16 % of pellet content, 30.12 % prey biomass). Another rodent prey identified was the common plantain squirrel, *Callosciurus notatus*, with 2 prey items (2.20% total, 4.45% biomass). Other prey identified in pellets were insects; grasshoppers (9.26% total and 0.11% biomass) and termites (12.65% total and 0.002% biomass).

In rice fields, a total of 90 pellets were collected with rodent pests being making up 99.52% of the prey biomass. The rice field rat, *Rattus argentiventer*, was the major diet of barn owls (96.77% total and 98.24% biomass) while shrews constituted a smaller fraction of the diet at 2.15% of total prey individuals and 1.28% of prey biomass. Amphibians were also recorded in the diet of barn owls in rice fields (1.07% total and 0.46% of prey biomass).

A total of 100 pellets were collected in oil palm plantations and 92.83% of total prey were rodents. The Malayan Wood Rat, *Rattus tiomanicus*, was the main prey species in terms of prey total (90.16%) and prey biomass (94.35%). Squirrels were also found in barn owl pellets with the diurnal rodent making up 2.45% of individual prey total and 3.71% of prey biomass. Grasshoppers were recorded as the second highest individual prey of barn owls in oil palm plantations (3.27%), though this group only make a small fraction of prey biomass (0.02%). A small percentage of the barn owl diet in oil palm plantations were made up of birds, reptiles and amphibians (4.07% total and 1.89% biomass).

The food niche breadth (FNB) of barn owls in all the areas was calculated to determine the dietary diversity of barn owls in each habitat (Table 1). The released barn owls in urban area recorded the highest FNB value at 2.90, indicating a high diet diversity of the introduced barn owls in the urban area. FNB value was second highest for barn owls in oil palm plantations (1.22 FNB) and barn owls in rice fields recorded the lowest FNB value (1.06 FNB).

### Prey weight of introduced urban barn owls

The biomass of identified preys inside the collected pellets were estimated using a standard log-log regression of right mandible length (mm) as a function body weight (g) as described by Hamilton (1980). Figure 2 shows the weight groups of introduced urban barn owl prey by numbers and biomass. Weights of prey were identified as extra small (< 3g), small (3-40g), medium (40-120 g) and large (120-160 g). Medium-sized prey were the most preferred weight group by owls (52.96% biomass and 38.71% total). Small-sized prey were the second highest preferred prey of introduced barn owls, making up 31.18% total prey consumed. However due to the small size, this prey category only contributed 16% of the prey biomass. Large-sized prey made up 12.90% of total prey and more than 30% of prey biomass. The extra small-sized prey made up 17.20% of total prey and contribute only 0.15% of prey biomass.

**Figure 2:**
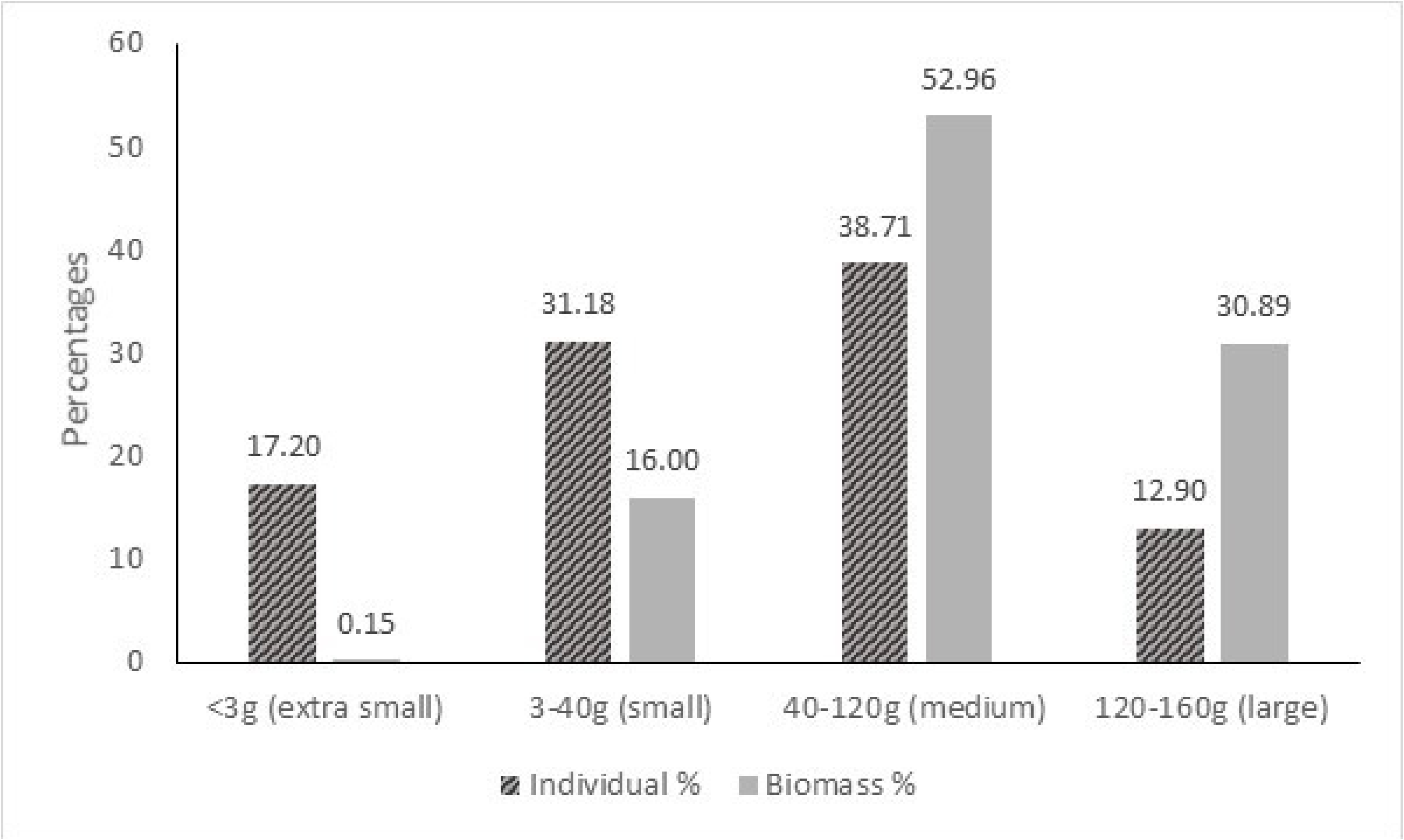
Percentages of individual and prey biomass of pellets of barn owls introduced in an urban areas.

As Norway rats were the most preferred prey of introduced barn owls in the urban area, further analysis was carried out on the size of the rats. Our analysis showed that the most consumed weight of rats were medium-sized rats, i.e. individuals weighing 80 to 100 g (Figure 3). Seventeen individual medium-sized rats were consumed (44.74%). Twelve small-sized rats weighing from 40 to 80 g were the second highest weight group consumed by barn owls (31.58%) and the less consumed weight group were large-sized rats weighing more than 120 g (9 individuals, 23.68%). Norway rats weighing less than 40 g and more than 160 g were not found in our pellet analysis.

**Figure 3:**
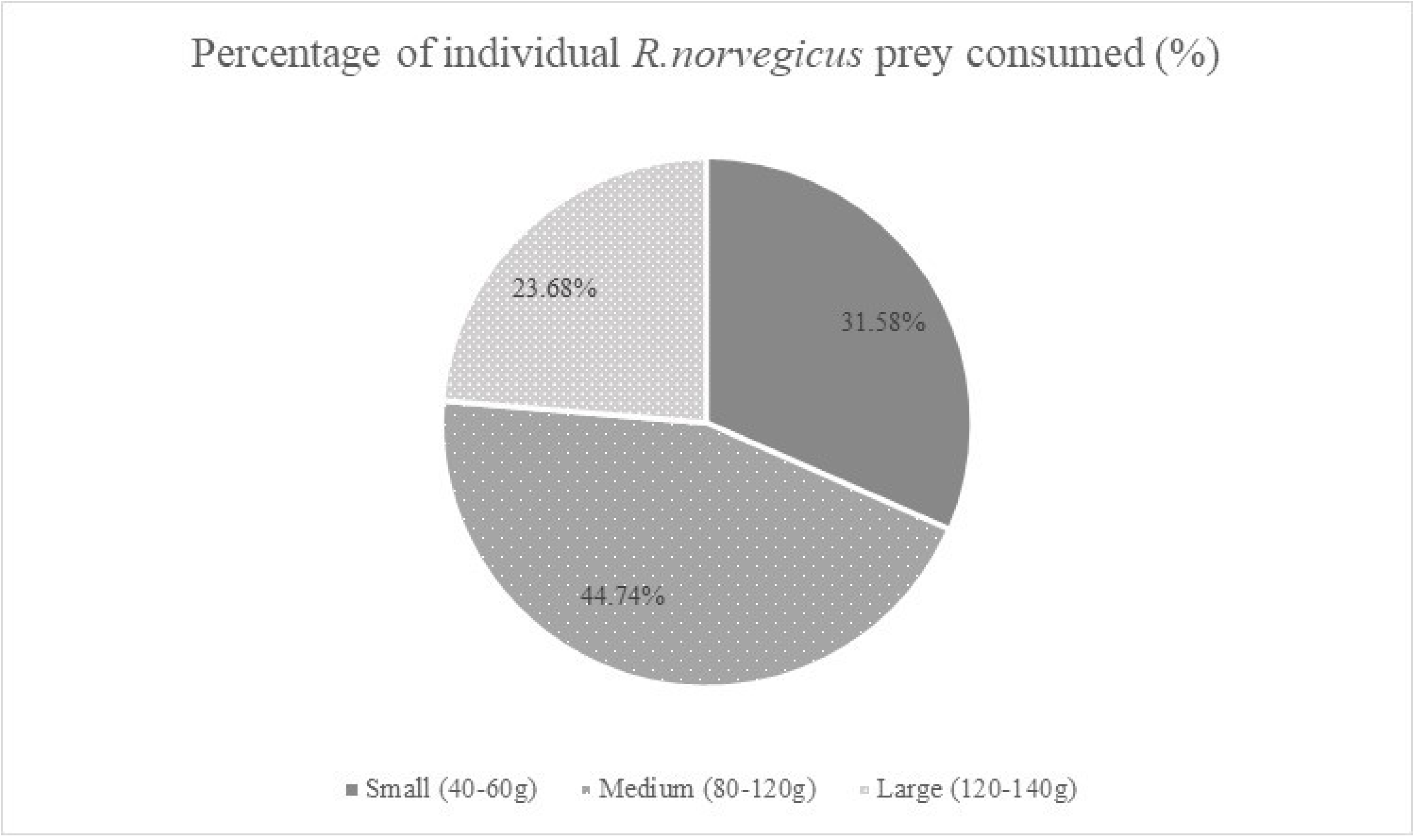
Percentage of size of Norway rats, R.norvegicus, prey consumed by barn owls introduced to an urban area.

## Discussion

### Diet of introduced barn owl and in agricultural landscapes

The barn owls in this study that were introduced and released in an urban area were seen roosting and perching in trees, roof spaces of buildings and houses, as well as abandoned structures. The owls were also seen hunting in open grass habitats near roadsides, human settlements, and backyards of shop lots. One of the barn owls also started occupying one of our installed nest boxes near the aviary, indicating the successful of release of barn owls in an urban area. On the other hand, a substantial amount of released barn owls in this study were untraceable a week after their release. These released young barn owls dispersed further away from the release site and are probably foraging around the urban areas of Penang Island or could have travelled further to mainland Peninsular Malaysia (Saufi et al., 2019).

Similar to various studies on the diet of barn owls, our study reports that barn owls in agricultural areas and urban areas prey mostly on small mammals. Norway rats and house shrews (80.12% total, 95.49% prey biomass) were the dominant prey group in the diet of barn owls in the urban area. Clark and Bunck (1991) reported that barn owls do consume commensal rodents along their distribution, though only in low frequencies. Our study shows that introduced barn owls were able to adapt to an urban setting and consume abundant urban rodent species. The high number of Norway rats and house shrew consumed by urban barn owls indicate the owls managed to hunt close to their release site and did not have to travel a great distance for more suitable open hunting grounds. Studies by Álvarez-Castañeda et al. (2004) in Mexico and Magrini and Facure (2008) in Brazil reported that pellets from barn owls in periurbans areas contain none to very little prey remnants from urban areas, suggesting that barn owls spend more time hunting in areas away from human settlements. In Canada, Hindmarch and Elliot (2015) reported that barn owls retained their preference for voles despite being in an urban landscape, although rats were consumed in higher amounts in urban areas.

The commensal rodent pests, Norway rats, *R. norvegicus*, and Black rats, *R. rattus*, are among the most widespread urban pest species in the world that resides frequently in close proximity to human habitation and are rarely found in the wild (Feng & Himsworth, 2014). The substantial occurrence of Norway rats inside the pellets of introduced barn owls show that the barn owls are taking advantage of the abundance of this pest species. Common house shrews were the second most consumed prey of barn owls in urban areas. These rodent species are the reported principal prey species in barn owl diet by several studies (Glue, 1967; Love et al., 2000; Hindmarch & Elliot, 2015; Horváth et al., 2018) and this species are more abundant in urban settings compared to agricultural settings (Chang et al., 1999). During tracking of released barn owls, Norway rats and house shrews were observed and frequently encountered in residential neighbourhoods, eateries, garbage dump areas and commercial areas within the study site (personal observation). It is however interesting to note that other detrimental rodent pests, i.e. house mice and roof rats, were not found in collected pellets despite being captured occasionally during rat trapping sessions as we conducted a study on population diversity of rats in urban areas around Penang Island. Timm (1994) documented that house mice and roof rats are typically found inside buildings and house mice rarely travel outside. Meanwhile, Norway rats and house shrews mainly inhabit and forage in open habitats (Timm, 1994), hence the two species inhabiting open areas and vegetation were the primary source of food for owls (Bonvicino & Bezerra, 2003).

Barn owls have been well documented to take advantage of other temporarily abundant types of prey that are vastly different from their usual diet, though extreme exceptions are unusual and usually occur in situations where small rodents are absence or scarce (Taylor, 1994). In Malaysia, most studies report that the diet of *T. alba javanica* is composed more than 90% of rats (Smal, 1990; Puan et al., 2011), with barn owls also preying on shrews, squirrels, birds and lizards in smaller numbers (Smal, 1990). Urban barn owls in this study consumed small rodents from the family *Sciuridae*. The common plantain squirrel, an uncommon barn owl prey, constituted a small fraction in the diet of urban barn owls (2.20% total, 4.45% biomass). An interesting result from our analysis is that there were no bird remnants found in the pellets of urban barn owls despite the abundant occurrence of passerine birds in our study site. In contrast, several reports analysing the diet of barn owls in rural and urban areas document that Norway rat, *R. norvegicus* and birds make up a high proportion of the diet of owls (Salvati et al., 2002; Teta et al., 2012; Hindmarch & Elliot, 2015). The pellet analysis of urban barn owls also showed that the owls preyed on insects, i.e. grasshoppers and termites. Though infrequent, barn owls have been reported to consume a high amount of insects, such as termites (e.g. Taylor, 1994) and locusts (e.g. Szabo et al., 2003; Shehab, 2005).

### Comparing diet of introduced urban barn owls and barn owls in agricultural areas

Though members of the Muridae family dominate the diet of barn owls in all habitats, the main prey species differed by habitat. *Rattus norvegicus* were the most preyed upon by introduced urban owls while *R.tiomanicus* and *R.argentiventer* were the most preyed upon small mammal in oil palm plantations and rice fields respectively. The barn owl prey-species preference in agricultural areas from our study is similar to other reports by Hafidzi and Naim (2003a) of barn owls in rice fields and Lenton (1984) of barn owls in oil palm plantations.

Food-niche breadth value of barn owls in the study was highest in urban areas compared to agricultural areas. There was a higher component of non-rodent prey items in urban areas compared to agricultural lands, with squirrels and insects accounting for 19.79% of individual prey and 4.51% of total prey biomass of owls in urban areas. This observation is similar to reports of Salvati et al. (2002) and Hindmarch et al. (2017) whom report an increased in non-rodent prey items in the pellets of barn owls as their habitat becomes more urbanized. Various food-niche studies showed barn owl prey selection was associated with rodent accumulations and responded to the density of rodents (e.g. Marti, 1988; Taylor, 1994; Leveau et al., 2006; Bernard et al., 2010; Marti, 2010; Milana et al., 2016). Similar to other food-niche analysis of barn owls in Europe (e.g., Milchev, 2015; Horváth et al., 2018), North America (Marti, 1988; 2010) and South America (e.g. Leveau et al., 2006; Teta et al., 2012), the low values of niche breadth analysis from agricultural areas in this study reflect the high abundance of an available and profitable prey, i.e. the dominance of *R.tiomanicus* and *R.argentiventer* in oil palm plantations and rice fields respectively. It is fairly well established that *R.argentiventer* is common in rice fields (Lam, 1983; 1988) and *R.tiomanicus* is common in oil palm plantations (Wood & Liau, 1984).

### Prey size preference of urban barn owls

Morphological features, such as body size and conspicuousness, and behaviour can also affect prey vulnerability to predation by barn owls (Derting & Cranford, 1989). Studies on differential prey selection by barn owls yield differing and often, contrasting results. Some studies show barn owls have an affinity to feed on smaller prey (e.g. Dickman et al., 1991; Rizuan *et al.*, 2017) while other studies have reported a tendency to feed on larger prey (Derting & Cranford, 1989; Castro & Jaksic, 1995). Our analysis show that barn owls prefer medium-sized Norway rats (40 to 120g) in urban areas, a finding similarly reported by Gaunt et al. (1997) and Hindmarch and Elliott (2015).

Barn owl diet also depends on the abundance of food supply, prey accessibility, which is affected by habitat characteristics, and general opportunistic feeding strategy (Taylor, 1994; Bond et al., 2005; Horváth et al., 2018; Arlettaz et al., 2010). As opportunistic predators, barn owls will hunt to maximize their nutrient intake and minimize energy expenditure (MacArthur & Pianka, 1966), hence prey size would play an important role in determining barn owl prey selection. Larger prey may be easy to locate but the energy gained might not compensate for the energy lost from subduing the prey (Ille, 1991), while smaller prey are hard to locate and more agile, hence energy gained might not compensate for the energy used to search and hunt for the prey (Colvin & McLean, 1986).

### Barn owls as urban rodent pest biological control agents

There are various ways to study the impact of released barn owls. Pellet analysis is a well-known and frequently used method to analyse owl prey content and preference (e.g., Bonvicino & Bezerra, 2003; Andrade et al., 2016). Meyer (2008) who studied the impact of released barn owls in a semi-urban area in Johannesburg evaluated the rodent population size using live-trapping before and after barn owl releases. Meyer (2008) reported a declining rat population following the release of barn owls. While these are positive reports, trap catchability could have biased the results as rats may developed trap shyness over time (Griffin, 2004). Additionally, some studies report that the mere presence of barn owls simply affects the behaviour of prey, i.e. the prey ventures less in the open (Abramsky et al., 1996).

Though some studies question the ability of barn owls to significantly reduce rodent populations (Van Vuren et al., 1998; Marti et al., 2005), results from this study show an affinity for barn owls to consume abundant commensal rodent pest species. While our results are preliminary, more studies are planned to further study the impact of introduced barn owls controlling rodent pest populations in an urban setting. Additionally, as commensal Norway rats are abundant and breed year-round, introduced urban barn owls would not have difficulty maintaining a high level of energy intake (Puan et al, 2011) and it is unlikely that the owls would switch prey species (Puan et al., 2011).

## Conclusion

Our study shows that barn owls in urban and agricultural areas are opportunistic predators that hunt almost exclusively on small mammal rats. Our study also showed that barn owls can adapt their prey species preference in different areas according to variations in small mammal abundance. Barn owls introduced in urban areas mostly consumed Norway rats and house shrews, which are notorious commensal rodent pests. Squirrels and insects were also preyed by these introduced urban barn owls but made up only a small fraction of their diet. Our results strongly indicate that barn owls introduced to urban areas have the potential to be an effective biological control agent against commensal rat pest populations following their high consumption by barn owls.

## Acknowledgements

We would like to thank Department of Agricultural, Bumbung Lima, Kepala Batas, Penang and Department of Agricultural Kerian District, Perak for giving permission to carry out our sampling in their rice fields. The authors also thank FGV Agri Services Sdn. Bhd and FGV R&D Sdn Bhd for providing the necessary facilities to conduct this study in their oil palm plantations. This work was supported by the Universiti Sains Malaysia Research University Grant (1001/PBIOLOGI/811270).

